# Flagellar genes are associated with the colonization persistence phenotype of the *Drosophila melanogaster* microbiota

**DOI:** 10.1101/2022.11.09.515865

**Authors:** Sarah J. Morgan, John M. Chaston

**Affiliations:** Department of Plant and Wildlife Sciences, Brigham Young University, Provo, Utah, USA

## Abstract

In this work we use *Drosophila melanogaster* as a model to identify bacterial genes necessary for bacteria to colonize their hosts independent of the bulk flow of diet. Early work on this model system established that dietary replenishment drives the composition of the *D. melanogaster* gut microbiota, and subsequent research has shown that some bacterial strains can stably colonize, or persist with, the fly independent of dietary replenishment. Here we reveal transposon insertions in specific bacterial genes that influence the bacterial colonization persistence phenotype by using a gene association approach. We initially established that different bacterial strains persist at varying levels, independent of dietary replenishment. We then repeated the analysis with an expanded panel of bacterial strains and performed a metagenome wide association (MGWA) to identify distinct bacterial genes that are significantly correlated with the colonization level of persistent bacterial strains. Based on the MGWA, we tested if 44 bacterial transposon insertion mutants from 6 gene categories affect bacterial persistence with the flies. We identified that transposon insertions in four flagellar genes, one urea carboxylase gene, one phosphatidyl inositol gene, one bacterial secretion gene, and one antimicrobial peptide (AMP) resistance gene each significantly influenced the colonization of an *Acetobacter fabarum* strain with *D. melanogaster*. Follow-up experiments revealed that each flagellar mutant was non-motile, even though the wild-type strain was motile. Taken together, these results reveal transposon insertions in specific bacterial genes, including motility genes, are necessary for at least one member of the fly microbiota to persistently colonize the fly.

**IMPORTANCE:** Despite the growing body of research on the microbiota, the mechanisms by which the microbiota colonizes a host can still be further elucidated. This study identifies bacterial genes that are associated with colonization persistence phenotype of the microbiota in *Drosophila melanogaster*, which reveals specific bacterial factors that influence establishment of the microbiota with its host. Identification of specific genes that affect persistence can help inform how the microbiota colonizes a host. Furthermore, a deeper understanding of the genetic mechanisms of the establishment of the microbiota could aid in further developing the *Drosophila* microbiota as a model for microbiome research.

## INTRODUCTION

*Drosophila melanogaster* is one of the best studied genetic models in existence. The genetics of *D. melanogaster* has been researched for over a century and the understanding of the genetics of *D. melanogaster* is thorough and expansive (1). One recent area of study that has gained attention in relation to *D. melanogaster* is that of microbiome studies. In *D. melanogaster* the microbiota can influence diverse *D. melanogaster* phenotypes and behaviors (2-4), including life history traits like fecundity (5), lifespan (6), and starvation resistance (7). The *D. melanogaster* gut microbiota resides primarily in the foregut and crop of the fly and is well characterized, usually comprising fewer than 100 bacterial taxa and numerically dominated by fewer than 10 taxa that are usually from the acetic acid bacteria, lactic acid bacteria, and enterobacteria (2, 8-10). Thus, when compared to vertebrates, many of which are dominated by hundreds of taxonomically diverse taxa (11, 12), the *D. melanogaster* microbiota is simpler. Also, the *D. melanogaster* microbiota is readily manipulated in lab conditions. Large numbers of *D. melanogaster* individuals can be made axenic, or mono-or or poly-associated with specific bacterial strains, enabling high reproducibility, and the techniques are relatively straightforward and have low financial costs (13). The expansive body of established research in *D. melanogaster* genetics and microbiota composition further assures ready access to genetic resources, strain collections and other tools to facilitate exploration of key biological questions (14).

The mechanisms and processes by which the microbiota colonizes *D. melanogaster* remain incompletely defined. Environmental bacteria gain access to the fly gut via horizontal transfer when the fly eats. An important early study suggested that the microbiota is established as flies ingest microbes in their diet, and that the microbial community is thereafter maintained by continuous consumption in the diet (15). Food travels through the entirety of the *D. melanogaster* gut in less than an hour (16), and the early work suggested that the microbiota was present as part of the bulk flow of food during this short transit time (14). At the same time, a separate study showed that the identity and abundance of the fly microbiota is inconstant within and across generations (17). Taken together, the primary conclusion was that the microbiota of *D. melanogaster* is transient and does not colonize the fly gut.

Later work studying the colonization of the *D. melanogaster* gut has refined this early view. It is now understood that some bacterial strains colonize their hosts and others do not, and that bacteria from wild flies generally colonize their hosts better than congeneric laboratory strains of bacteria. Some of these works show that bacterial isolates can proliferate in the fly gut, allowing for stable association with the host independent of continuous uptake through diet (2, 18, 19).Specific bacterial genes that might responsible for colonization processes by were suggested by an analysis that showed uric acid degradation and flagellar genes are primarily present in bacteria isolated from wild, but not laboratory, *D. melanogaster* lines (20). The main location of bacterial colonization is likely the foregut because this gut region that bears the highest bacterial loads, and bacterial colonization is readily visualized within this structure (2, 8, 10). Additionally, analyses of the microbiomes of flies and their diets showed that the bacterial identities and abundances in fly diet do not necessarily reflect the bacterial content and abundance of the host (21). Taken together, these more recent findings have suggested that host factors are not solely responsible for the establishment on the microbiota in the host, and that microbial factors also play key but incompletely defined roles in the process of host colonization.

The goal of this study was to identify microbial factors that affect bacterial persistence in the host, *D. melanogaster*. To test if specific bacterial genes influence bacterial persistence with the host we asked 3 questions: 1) Is there a significant difference in bacterial persistence levels between bacterial strains? 2) Can differences in bacterial persistence be correlated with the presence of specific bacterial genes in the colonizing or non-colonizing bacterial strains? 3) Can a mutant analysis confirm the associations predicted in step 2? Using a high throughput assay involving serial transfers of flies onto sterile food, a metagenome-wide association, and functional analysis of bacterial transposon insertion mutants, we identified transposon insertions in four flagellar genes, one urea carboxylase gene, one phosphatidyl inositol gene, one bacterial secretion gene, and one antimicrobial peptide (AMP) resistance gene that each significantly affected the ability of *Acetobacter fabarum* to persist with *D. melanogaster*. These transposon insertions implicate specific microbial factors as necessary for bacteria to persist with *D. melanogaster*.

## RESULTS

### There is species-specific variation in bacterial persistence with the host

To begin, we developed a high throughput assay to test the colonization persistence phenotype of a taxonomically diverse panel of bacterial strains in the flies. Our goal was to identify a regime of serial transfers that would allow us to measure bacterial persistence across a range from highly proficient to completely deficient. We detected a wide range of CFU abundances in flies that were colonized from birth with a preliminary panel of seven bacterial strains and then, when 3 day-old-adults, were serially transferred to sterile diets six times total over two days (Fig. 1, Fig. S1, Table S1). The abundances of bacteria in male and female flies reared with the seven bacterial strains from three different families varied in their abundance in the flies from 0 to > 88,000 CFU fly^-1^ (Kruskal Wallis (KW) χ^2^_13,380_ = 191.09, p < 1 × 10^−15^). There were no significant differences between the CFU abundances in male and female flies (Fig. 1), a somewhat unexpected finding because female flies usually bear higher bacterial loads than male flies. The finding that male and female flies bore comparable bacterial loads following frequent transfer to sterile diets suggests that the persistent microorganisms may occupy similar spaces and niches between the two sexes, at least for these tested strains. We focused on just one sex in our subsequent assays and elected to study female flies, in part to make our results comparable with other analyses that measured life history traits of mono-associated females flies (5-7).Overall, these results confirm that our assay allowed us to detect a range of persistence phenotypes by different bacterial strains that did not vary significantly with the sex of the flies.

**Figure 1.**
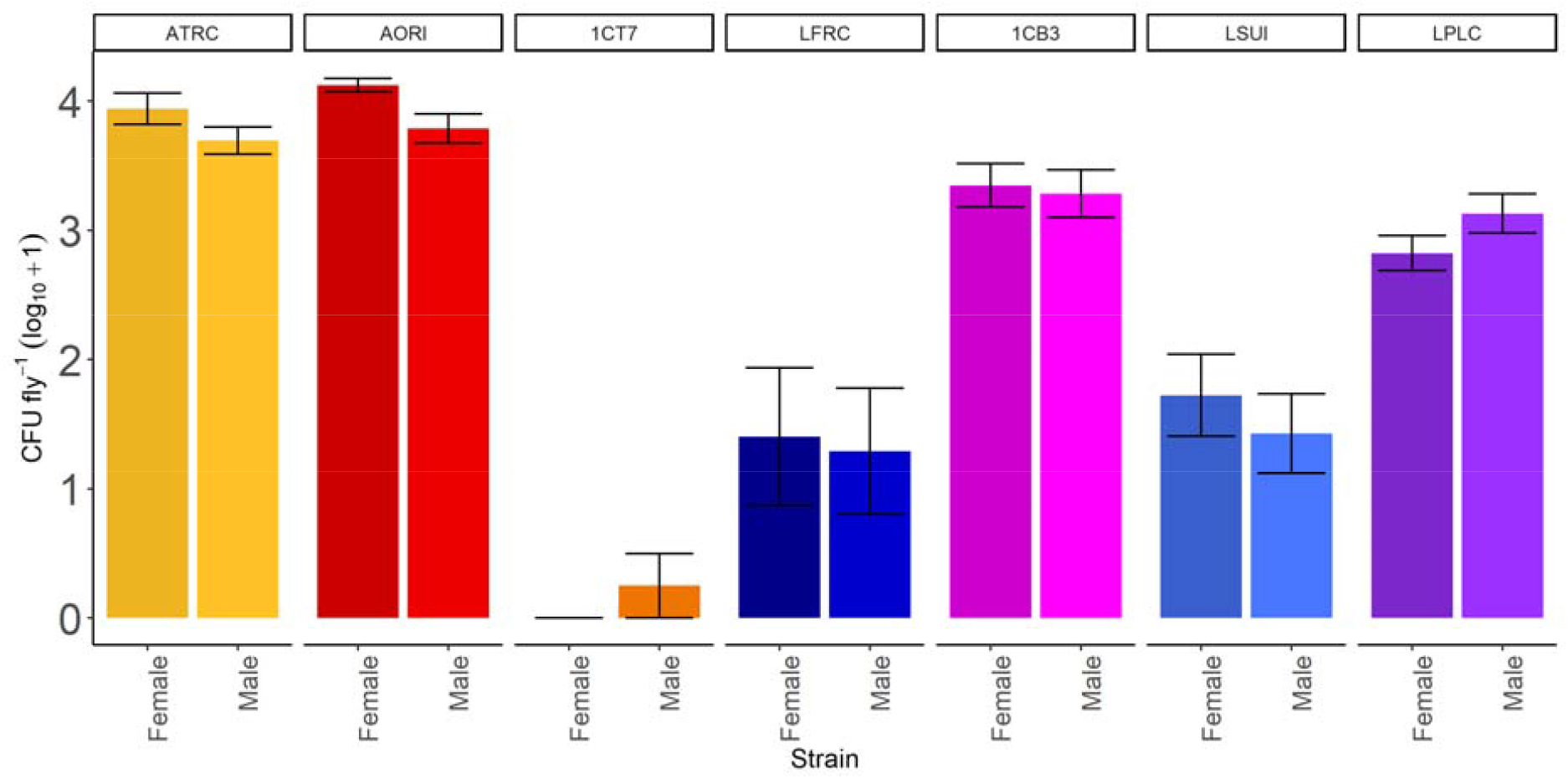
Bacterial persistence with the flies is strain-specific. Significant differences between the sexes were determined by a Wilcoxon test. Table 3 reports the strain names of the 4-character codes.

### Flagellar motility is among the pathways significantly associated with bacterial persistence

To predict bacterial genes that contribute to bacterial persistence in the flies, we measured CFU loads of 41 different bacterial strains in the flies after six serial transfers and statistically associated the bacterial loads with bacterial gene presence-absence patterns. The 41 different strains showed a wide range of CFU abundances in the flies (KW)χ^2^ _6,380_ = 176.04, p < 1 × 10^−15^), providing excellent strain-level phenotypic variation (Fig 2, Fig S2, Table S2). Then, we performed an MGWA to identify bacterial genes whose presence was associated with this variation in CFU counts. We measured the association between bacterial persistence and 12,105 orthologous groups (OGs) that were collectively spread across 3,760 phylogenetic distribution groups (PDGs, a unique set of taxa in which an OG is present). We determined that the presence-absence patterns of 385 Ogs were statistically associated with bacterial persistence in the flies (Bonferroni-corrected p < 0.01).

**Figure 2.**
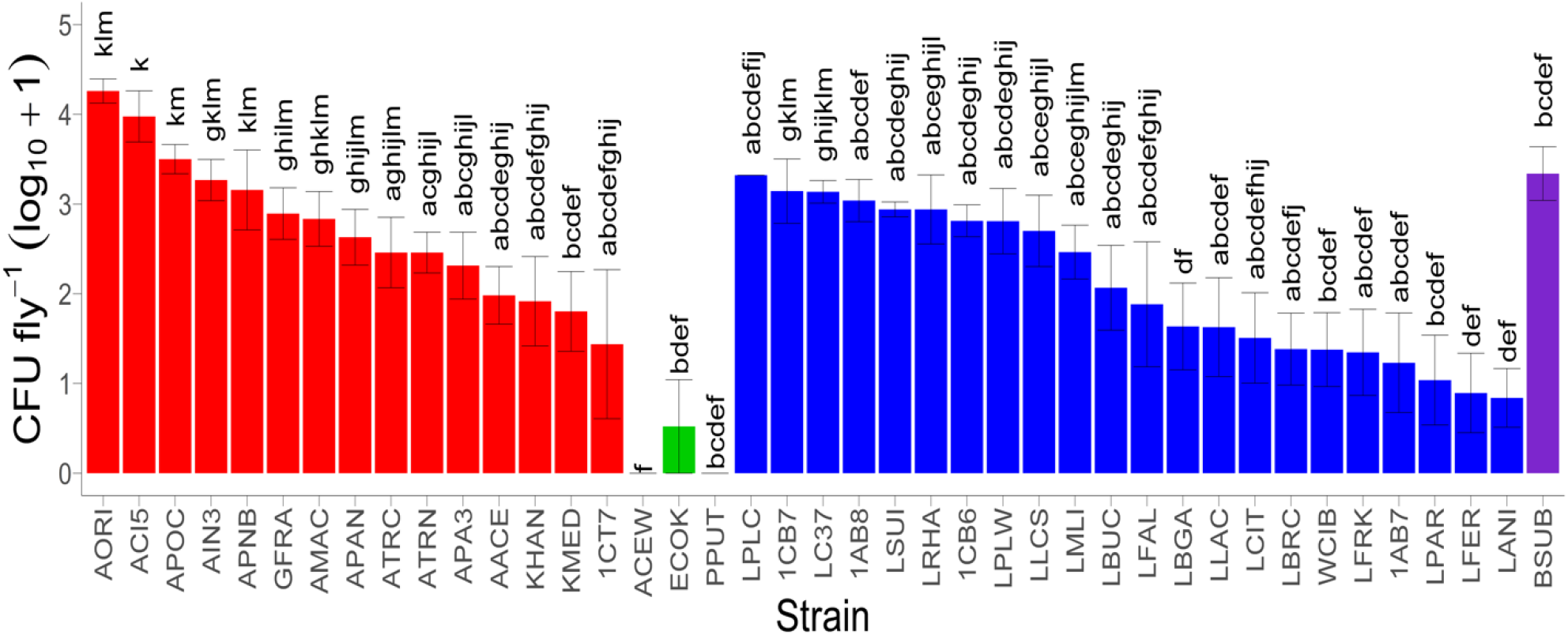
Strain-specific differences in persistence of a broad panel of bacteria. Shading matches bacterial groups: acetic acid bacteria (red), gammaproteobacteria (green), lactic acid bacteria (blue), non-lactic acid firmicutes (purple). Different letters above each bar show significant differences in bacterial persistence between strains by a Kruskal-Wallis test followed by a post-hoc Dunn test. Table 3 reports the strain names of the 4-character codes.

From the OGs that were significantly associated with changes in bacterial persistence, we selected a subset to focus on in a mutant analysis. We chose mutants that we could obtain from an existing library of mapped and arrayed *A. fabarum* transposon insertion mutants (22), and that corresponded to genes in pathways that were enriched among the top MGWA hits. We assigned the 385 significant KEGG IDs from the MGWA to KEGG pathways and performed an enrichment analysis that revealed the membership of the top KEGG matches were enriched for 10 KEGG pathways (Table 1, complete results in Table S4). Of these, we selected to test 26 different genes from 4 KEGG pathways for further analysis: phosphatidyl inositol signaling system, bacterial secretion system, nicotinate and nicotinamide metabolism, and cationic antimicrobial peptide (CAMP) resistance. Among the bacterial secretion system genes were genes encoding for the flagellar apparatus, which was consistent with a previous suggestion that flagellar genes were likely involved in bacterial persistence with the flies (20). From these candidates we selected 44 mutants from five functional pathways for follow-up mutant analysis (Table 2).

**Table 1.** Significant results in KEGG pathway enrichment analysis

| Pathway | Map ID | P-Value |
| --- | --- | --- |
| *Staphylococcus aureus* infection | map05150 | 2.22E-06 |
| Biofilm formation - *Pseudomonas aeruginosa* | map02025 | 4.11E-06 |
| D-Alanine metabolism | map00473 | 4.92E-05 |
| Nicotinate and nicotinamide metabolism | map00760 | 0.0005 |
| Bacterial secretion system | map03070 | 0.002 |
| Phosphatidylinositol signaling system | map04070 | 0.03 |
| Atrazine degradation | map00791 | 0.03 |
| Cationic antimicrobial peptide | map01503 | 0.044 |
| Vancomycin resistance | map01502 | 0.046 |
| Arginine biosynthesis | map00220 | 0.0499 |

**Table 2.** Bacterial mutants used in this study.

| Code | Gene | Kegg | P-Value |
| --- | --- | --- | --- |
| Urea Carboxylase |  |  |  |
| UCA1<sup>†</sup> | E6.3.4.6; urea carboxylase [EC:6.3.4.6] | K01941 | < 10<sup>-5</sup> |
| UCA2*<sup>†</sup> | E6.3.4.6; urea carboxylase [EC:6.3.4.6] | K01941 | < 10<sup>-5</sup> |
| Phosphatidylinositol Signaling |  |  |  |
| suhB1 | E3.1.3.25; myo-inositol-1(or 4)-monophosphatase [EC:3.1.3.25] | K01092 | 0.002 |
| suhB2* | E3.1.3.25; myo-inositol-1(or 4)-monophosphatase [EC:3.1.3.25] | K01092 | 0.002 |
| Bacterial Secretion |  |  |  |
| secB* | 6666666.222946.peg.857 *secB*; preprotein translocase subunit SecB | K03071 | 0.04 |
| yajC1 | *yajC*; preprotein translocase subunit YajC | K03210 | 0.02 |
| yajC2 | *yajC*; preprotein translocase subunit YajC | K03210 | 0.02 |
| tolC1 | *tolC*; outer membrane protein | K12340 | 0.02 |
| tolC2 | *tolC*; outer membrane protein | K12340 | 0.02 |
| Nicotinic Acid |  |  |  |
| pncB1 | pncB; nicotinate phosphoribosyltransferase [EC:6.3.4.21] | K00763 | 0.02 |
| pncB2 | pncB; nicotinate phosphoribosyltransferase [EC:6.3.4.21] | K00763 | 0.02 |
| pncC1 | *pncC*; nicotinamide-nucleotide amidase [EC:3.5.1.42] | K03743 | 1 |
| pncC2 | *pncC*; nicotinamide-nucleotide amidase [EC:3.5.1.42] | K03743 | 1 |
| pnuC1 | nicotinamide transporter, PnuC-like | K03811 | < 10<sup>-5</sup> |
| pnuC2 | nicotinamide transporter, PnuC-like | K03811 | < 10<sup>-5</sup> |
| nadC1 | nadC; nicotinate-nucleotide pyrophosphorylase (carboxylating) [EC:2.4.2.19] | K00767 | 0.02 |
| nadC2 | nadC; nicotinate-nucleotide pyrophosphorylase (carboxylating) [EC:2.4.2.19] | K00767 | 0.02 |
| nadD | *nadD*; nicotinate-nucleotide adenyltransferase [EC:2.7.7.18] | K00969 | 1 |
| AMP Resistance |  |  |  |
| degP1 | *degP*; serine protease Do [EC:3.4.21.107] | K04771 | 0.02 |
| degP2 | *degP*; serine protease Do [EC:3.4.21.107] | K04771 | 0.02 |
| degP3 | *degP*; serine protease Do [EC:3.4.21.107] | K04771 | 0.02 |
| degP4 | *degP*; serine protease Do [EC:3.4.21.107] | K04771 | 0.02 |
| degP5 | *degP*; serine protease Do [EC:3.4.21.107] | K04771 | 0.02 |
| degP6 | *degP*; serine protease Do [EC:3.4.21.107] | K04771 | 0.02 |
| mprF1* | *mprF*; phosphatidylglycerol lysyltransferase [EC:2.3.2.3] | K14205 | 0.04 |
| mprF2 | *mprF*; phosphatidylglycerol lysyltransferase [EC:2.3.2.3] | K14205 | 0.04 |
| amiABC1 | N-acetylmuramoyl-L-alanine amidase [EC:3.5.1.28] | K01448 | < 10<sup>-5</sup> |
| amiABC2 | N-acetylmuramoyl-L-alanine amidase [EC:3.5.1.28] | K01448 | < 10<sup>-5</sup> |
| Flagellar Assembly |  |  |  |
| fliP | *fliP*; flagellar biosynthetic protein FliP | K02419 | 0.42 |
| fliM1 | *fliM*; flagellar motor switch protein FliM | K02416 | 1 |
| fliM2 | *fliM*; flagellar motor switch protein FliM | K02416 | 1 |
| fliH1 | *fliH*; flagellar assembly protein FliH | K02411 | 1 |
| fliH2 | *fliH*; flagellar assembly protein FliH | K02411 | 1 |
| fliG1 | *fliG*; flagellar motor switch protein FliG | K02410 | 1 |
| fliG2 | *fliG*; flagellar motor switch protein FliG | K02410 | 1 |
| fliF1*<sup>†</sup> | *fliF*; flagellar M-ring protein FliF | K02409 | 1 |
| fliF2*<sup>†</sup> | *fliF*; flagellar M-ring protein FliF | K02409 | 1 |
| fliI1<sup>†</sup> | *fliI*; flagellum-specific ATP synthase [EC:3.6.3.14] | K02412 | 1 |
| fliI2*<sup>†</sup> | *fliI*; flagellum-specific ATP synthase [EC:3.6.3.14] | K02412 | 1 |
| flgK1 | *flgK*; flagellar hook-associated protein 1 FlgK | K02396 | 1 |
| flgK2 | *flgK*; flagellar hook-associated protein 1 FlgK | K02396 | 1 |
| flgE1*<sup>†</sup> | *flgE*; flagellar hook protein FlgE | K02390 | 1 |
| flgE2<sup>†</sup> | *flgE*; flagellar hook protein FlgE | K02390 | 1 |
| flgF1 | *flgF*; flagellar basal-body rod protein FlgF | K02391 | 1 |
| flgF2 | *flgF*; flagellar basal-body rod protein FlgF | K02391 | 1 |
| flgH1*<sup>†</sup> | *flgH*; flagellar L-ring protein precursor FlgH | K02393 | 1 |
| flgH2*<sup>†</sup> | *flgH*; flagellar L-ring protein precursor FlgH | K02393 | 1 |
| flgI1 | *flgI*; flagellar P-ring protein precursor FlgI | K02394 | 1 |
| flgI2 | *flgI*; flagellar P-ring protein precursor FlgI | K02394 | 1 |
| flgB | *flgB*; flagellar basal-body rod protein FlgB | K02387 | 1 |
| flhB1 | *flhB*; flagellar biosynthetic protein FlhB | K02401 | 1 |
| flhB2 | *flhB*; flagellar biosynthetic protein FlhB | K02401 | 1 |
| flgA1 | *flgA*; flagella basal body P-ring formation protein FlgA | K02386 | 1 |
| flgA2 | *flgA*; flagella basal body P-ring formation protein FlgA | K02386 | 1 |

<sup>†</sup> Bonferroni corrected p-value from the MGWA.
\* significantly difference persistence phenotype from the wild-type
<sup>†</sup> tested for motility.

### Flagellar motility genes are among those that influence bacterial persistence with the flies

We measured the persistence phenotype of bacterial mutants for genes identified in the MGWA as a step towards validating MGWA predictions and identifying bacterial gene candidates that influence bacterial persistence with the flies. We identified 10 transposon insertions mutants in 8 different genes from 5 different gene categories that significantly influenced bacterial persistence with the flies (Fig. 3, Table S5; KWχ^2^_56,1239_ = 189.32, p < 10^−15^). Nine of the mutants conferred a lower persistence phenotype, and one mutant, myo-inositol-monophosphatase, conferred a higher persistence phenotype. Six of the significant mutants were flagellar assembly mutants, representing four flagellar genes. These efforts confirm that multiple different bacterial pathways influence bacterial persistence with the flies, and especially implicate bacterial flagellar genes as possible effectors of this phenotype.

**Figure 3.**
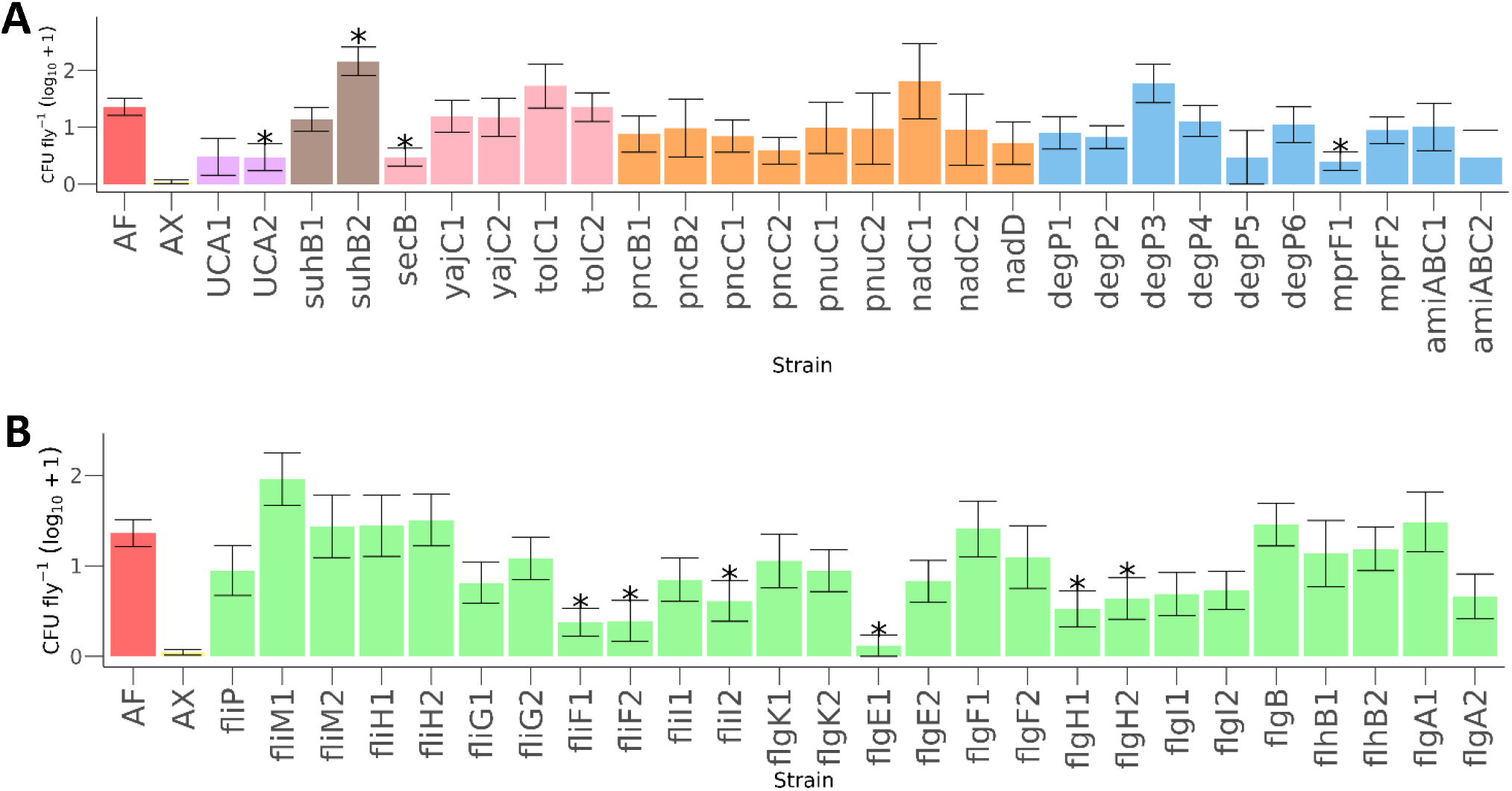
Transposon insertion mutants affect the bacterial persistence phenotype. Asterisks indicate that the bacterial mutants had a significantly different persistence phenotype from the wild-type bacterial strain (*A. fabarum*, AF), determined by pairwise Krukal-Wallis tests (p< 0.05). (A) All non-flagellar mutants testing in the assay. Shading matches type of mutants tested: wild-type *A. fabarum* (red), bacteria-free (AX) flies (yellow), urea carboxylase (purple), phosphatidylinositol (brown), bacterial secretion (pink), nicotinate metabolism (orange), AMP resistance (blue). (B) All flagellar mutants tested in the assay, shown in green.

### Flagellar mutants that persist poorly with the flies are non-motile

To determine if bacterial motility is associated with bacterial persistence, we compared the motility phenotypes of flagellar mutants, non-flagellar mutants, and wild-type *A. fabarum*. All flagellar mutants that were tested were non-motile, while the wild-type *A. fabarum* and the non-flagellar mutants were all motile (Fig. 4, Table S6; ANOVA, linear mixed-model, p < 5.176 × 10^−15^, f-value = 13.662). The correlation between flagellar mutants involved in bacterial persistence and non-motility indicates that bacteria being able to move may play a role in bacterial persistence in flies.

**Figure 4.**
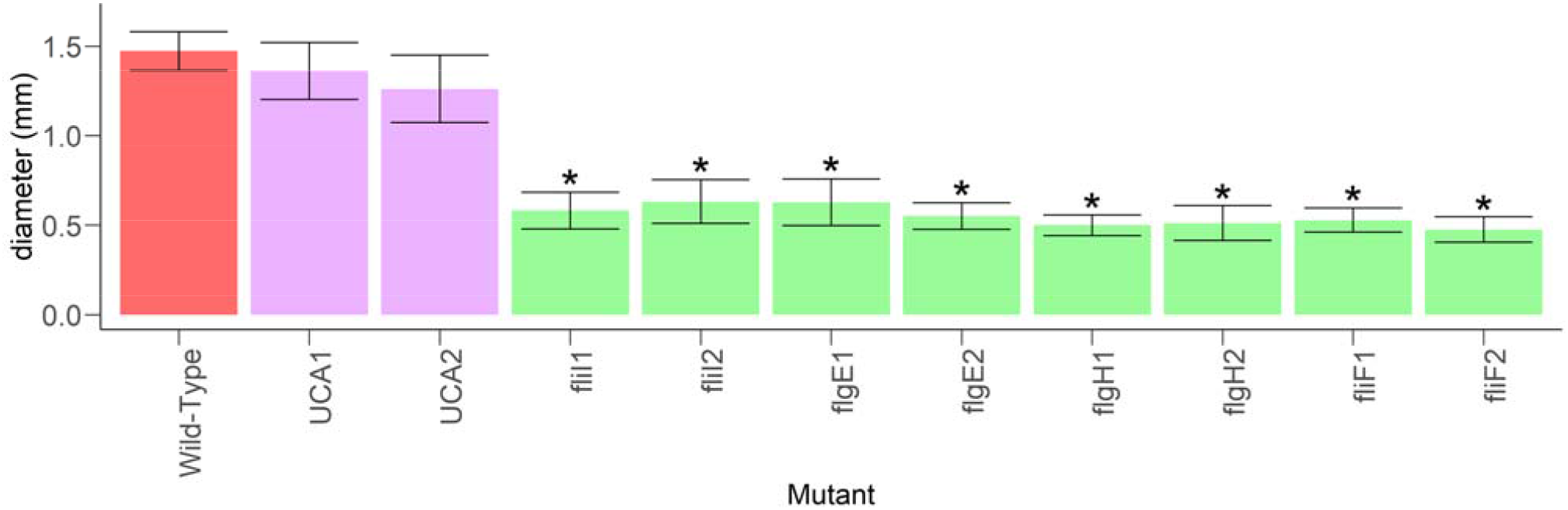
Flagellar mutants that influence persistence are non-motile. The mean diameter measured on the halos of the motility tests for each bacterial strain are shown. Shading matches type of mutants tested: wild-type *A. fabarum* (red), urea carboxylase (purple), flagellar assembly (green). Asterisks are shown if the average diameter of a mutant strain differed significantly from that of the wild-type strain. Significant differences between diameters were determined by a linear mixed-effects model with a post-hoc Dunnett test applied.

## DISCUSSION

This work adds to the growing knowledge of how the microbiota is established and persists with *D. melanogaster*. Identifying bacterial genes that are involved in bacterial persistence with the flies shows that the host alone is not responsible for the establishment and persistence of the gut microbiota. In particular, we identified that transposon insertions in four flagellar genes, one urea carboxylase gene, one phosphatidyl inositol gene, one bacterial secretion gene, and one AMP resistance gene affect bacterial persistence of a representative member of the gut microbiota, *A. fabarum*. These findings identify candidate pathways and mechanisms that underlie bacterial persistence of the microbiota.

Here we reveal that transposon insertions in four flagellar genes affect bacterial persistence with the flies. Motility is associated with the function of these flagellar genes because each persistence-deficient flagellar mutants was also non-motile, suggesting that motility contributes to bacterial persistence in the flies. We have not defined any specific roles for motility in bacterial persistence with the flies, but a possible explanation could be that motility enables bacteria to find favorable niches through bacterial taxis within the *Drosophila* foregut. A favorable niche could be a region of the fly that has better quantity or quality of nutrients, less immune stress, or decreased competition from other microorganisms. Also, other studies have shown that bacteria with intact, motile flagella are at an advantage for host colonization (23-25). Alternatively, these flagellar genes may be involved in bacterial secretion (26). Bacterial secretion could create a favorable environment or be beneficial for the host. Bacterial surface components, including flagella, are also known to interact directly with the gut epithelium, which could also promote persistence (27). Counter to either of these expectations is the finding that some high-persisting bacterial strains like *Acetobacter pomorum* DmCS_004 lack flagellar motility genes (Table S4). Therefore, if motility is necessary for colonization then *A. pomorum* must do so independent of prototypical flagellar machinery; or, alternatively, motility may be one of several redundant mechanisms that contribute to bacterial persistence with the host.

In addition to flagellar motility genes, we identified transposon insertions in other genes that were significantly associated with variation in bacterial persistence with the flies. One of these is in an AMP resistance gene, the flippase *mprF*, which translocates lysyl-phosphatidylglycerol to the outer leaflet of the membrane. It is also involved in resistance to multiple antimicrobial peptides from the host and other competing microorganisms (28). AMP resistance has already been identified as a mechanism for stable association of gut commensals in the human gut during periods of host inflammation (29). Pathogens are also able to stably colonize the gut due to AMP resistance (30), so it is possible that beneficial microbes also stably colonize the gut through AMP resistance. Some AMP resistance genes, such as *degP* (although not significant in our analysis) have been shown to help bacteria adjust to survive at high temperatures by decreasing temperature-sensitive growth (31). AMP resistance in bacteria may be important because AMPs were shown to be necessary in *D. melanogaster* to regulate microbiota abundance in the gut (32). Overall, AMP resistance can help microbes to colonize the gut by protecting against host antimicrobial peptides and helping the bacteria to adjust to a changing environment.

Of 3 bacterial secretion genes we tested, only transposon insertions in *secB*, which is involved in the quorum sensing, protein export, and bacterial secretion system pathways, significantly affected bacterial persistence with the flies. *SecB* specifically exports proteins, but is also involved in stress-responsive type II toxin-antitoxin (TA) systems (33) which have been shown to be important in niche-specific colonization of *Escherichia coli* in humans (34). Another gene, *yajC*, is also involved in these pathways but not shown to be significant. Identification of some, but not all genes in a pathway as significant for bacterial persistence leads to the idea that only particular parts of the pathways are important for the bacterial persistence phenotype. The sec bacterial secretion system is used in pathogenic bacteria to secrete virulence factors (35), hinting at a possible interaction with the host. Bacterial secretion can also aid the bacteria in establishing a hospitable niche by secreting products such as AMP resistance products which are known substrates of SecB (36).

The other two transposon insertions that significantly affected bacterial persistence with the flies were in urea carboxylase and myo-inositol-1(-or 4)-monophosphatase genes. Urea carboxylase is involved in arginine biosynthesis, atrazine degradation, and metabolic pathways. Myo-inositol-1(-or 4)-monophosphatase is involved in streptomycin biosynthesis, inositol phosphate metabolism, metabolic pathways, biosynthesis of secondary metabolites, and phosphatidylinositol signaling system. Since these genes are not closely related to other significant genes and are involved in many different pathways, further research would be required to understand the role they play in bacterial persistence.

### Future directions

In conclusion, we show here that transposon insertions in bacterial genes, particularly flagellar genes, are implicated in bacterial persistence in flies. Further studies could test more genes, more bacteria, and different hosts to identify genes that were not tested here and to see if the genes associated with persistence in this study are fly specific. Furthermore, the further characterization of the genes we identify here could provide insight into their roles in bacterial persistence.

Additionally, hypothetical proteins could also be studied to determine if undiscovered proteins play a role in persistence. Together, additional such approaches will add to the growing body of literature explaining how the fly microbiota specifically is established, and, more generally, how bacteria colonize their hosts.

## MATERIALS AND METHODS

### Bacterial and fly cultures

The fly stock was originally obtained from Mariana Wolfner at Cornell University and is a Wolbachia-free stock of Canton-S *D. melanogaster* flies. The stock flies were raised in an incubator on a 12-h light-dark cycle at 25°C. They were raised on a yeast-glucose (YG) diet that contains 10% brewer’s yeast, 10% glucose, 1% agar, 0.084% propionic acid, and 0.08% phosphoric acid.

Stocks of bacterial strains were stored at -80°C. The bacterial strains were streaked for isolation onto clade-specific media plates and incubated at 30°C for 2-3 days (Table 3). The different media types were: mMRS (Criterion C5932), LB (Apex 11-119), and potato dextrose (Sigma-Aldrich 70139-500G). Aerobic strains were placed in the incubator while anaerobic strains were put in carbon dioxide-flooded containers that were sealed and put in the incubator. One colony was then removed from the plates and placed in a tube of 5mL of clade-specific media broth and incubated at 30°C for 1-2 days. If the strains were aerobic the tubes of liquid broth were grown under oxic conditions by shaking. Aerotolerant strains were raised under microoxic conditions by remaining static. The bacteria were then diluted in a 1:8 dilution four times and normalized to OD_600_ of .01.

**Table 3.**
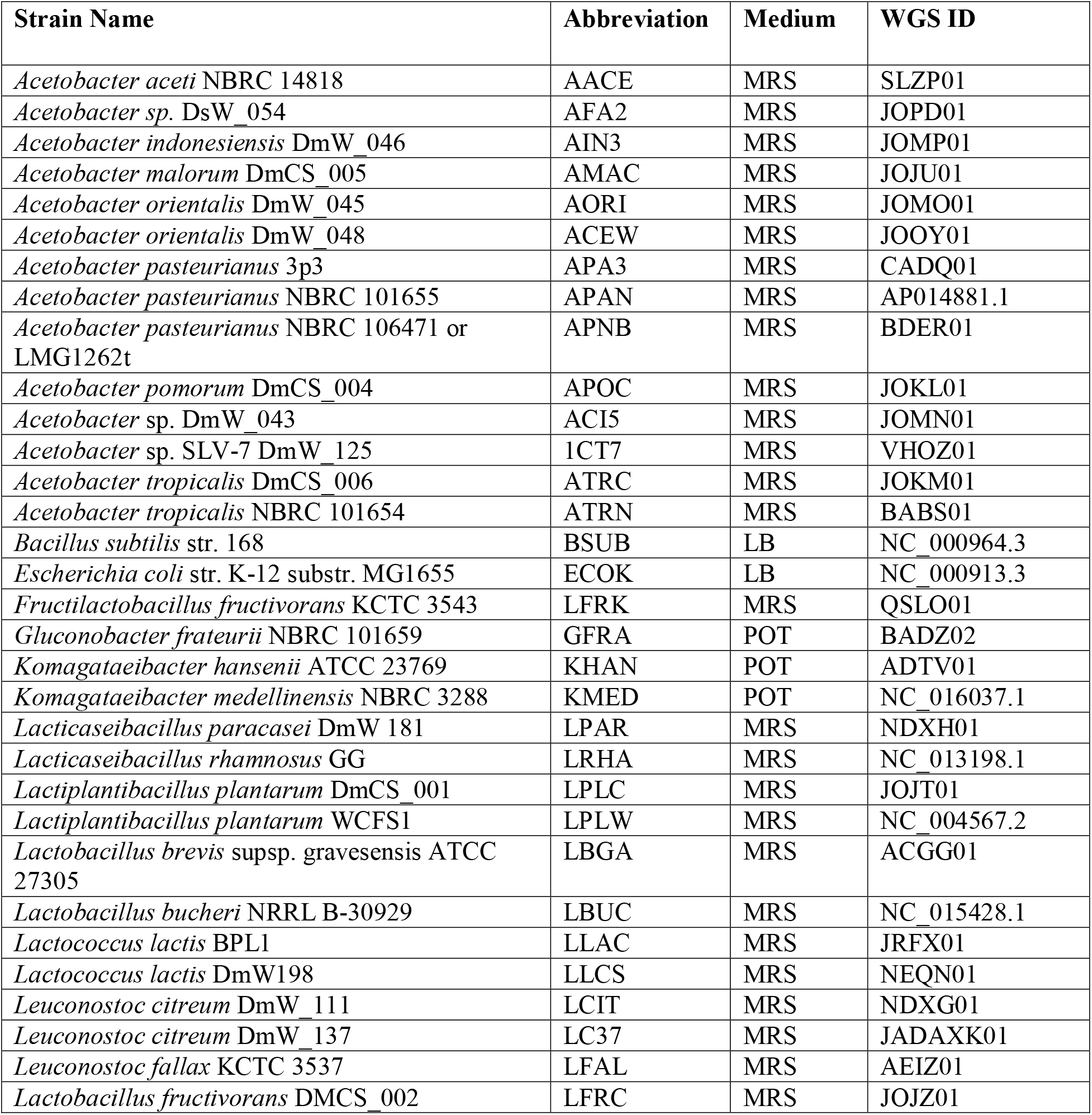
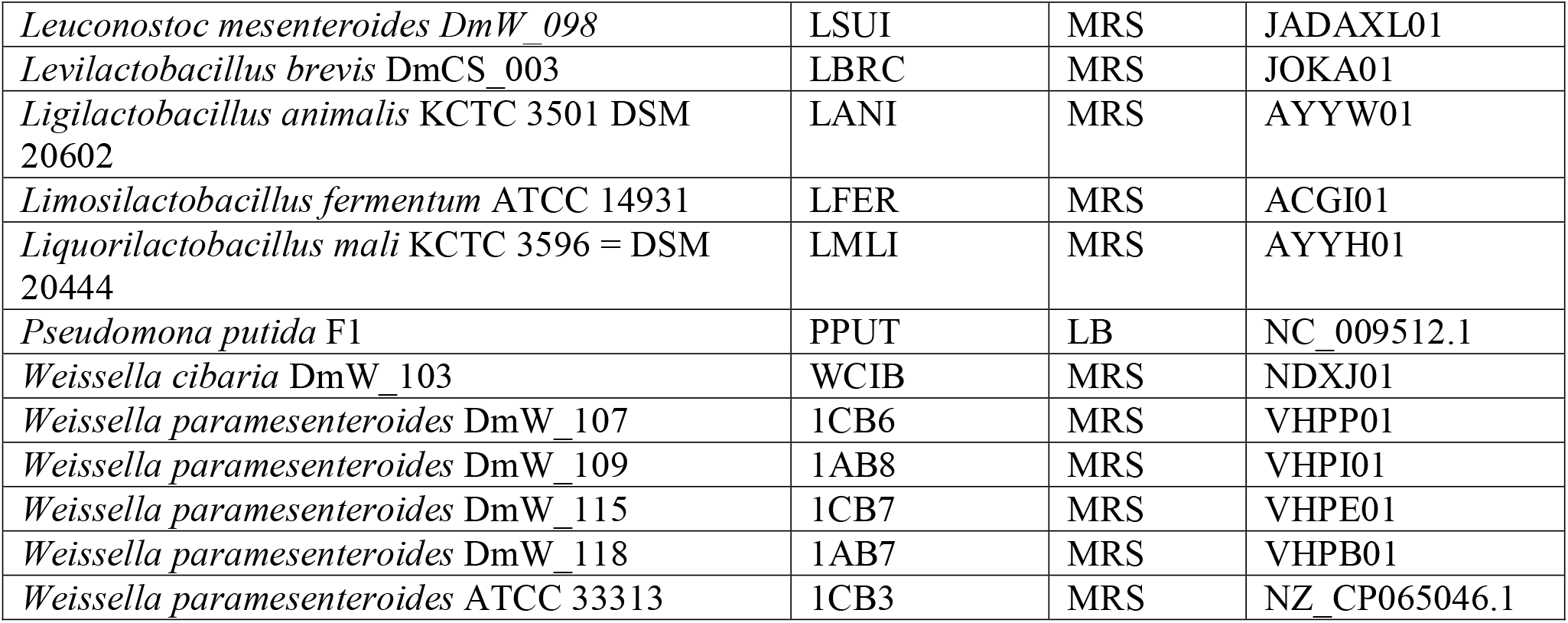
Bacterial strains used in this study

### Axenic and mono-associated flies

All flies used in the persistence assay were derived as bacteria-free embryos before they were inoculated with bacteria. Fly eggs were made axenic by removing the chorion layer of eggs. To do this, stock flies were allowed to lay eggs for 18-20 hours on a plate made of 10% brewer’s yeast, 10% glucose, 1% agar, and grape juice. The eggs were then collected and washed with a 0.6% hypochlorite solution twice for 2.5 minutes each. They were then washed three times with double distilled, autoclaved water. Then, 40-60 eggs were transferred into 50 mL vials containing 7.5 mL of autoclaved YG diet.

To mono-associate the flies, 50 μL of normalized bacteria were inoculated to axenic eggs in the sterile diet. The fly vials were then placed in a tray and put in an incubator at 25 ° C with a 12-hour light-dark cycle.

### Persistence assay

Bacterial persistence with the flies was measured using an assay that frequently transferred adult flies to a sterile diet. Four days post bulk eclosion of the flies, 4 female flies from each vial were transferred under carbon dioxide anesthesia into separate wells of a 96 well plate with 150 μL of sterile diet at the bottom. The flies were then transferred to new 96-well plates containing sterile food 3 times a day (8AM, 1PM, 6PM) for 2 days. After the last transfer, the flies were placed in 1.7 mL microcentrifuge tubes with 150 μL of PBS and 150 μL of ceramic beads and homogenized in a GenoGrinder for 2 minutes at 1750rpm. The contents of the microcentrifuge tubes were then dilution plated and cultured in an incubator at 30°C until colonies were large enough to count (around 2-3 days), each colony was counted as one colony forming unity (CFU) and used as a measure of persistence. If the colonies were too dense to count, then 160 was used as the CFU number, corresponding to 128,000 CFU fly^-1^). The first analysis was performed with 7 strains, and a Kruskal-Wallis test and pairwise Wilcoxon tests between sexes for each strain were performed to assess if the CFU per fly were significantly different. The second analysis was performed with 41 different strains for use in an MGWA, and a Kruskal-Wallis test with post-hoc all-against-all-pairwise Wilcoxon tests were performed to determine significance groups of CFU per fly between all strains. The third analysis was performed with 44 mutants identified through the MGWA, and a Kruskal-Wallis test was performed to test if each mutant was significantly different from the wild-type control. In each experiment, each treatment had triplicate vials in each of three separate experiments. Vials were discarded from the analysis if they were contaminated or the vial density was less than 30 flies. A vial was determined to be contaminated if undiluted aliquots bore more than 5 CFU of an unexpected colony morphology.

### Metagenome-wide association

A metagenome-wide association (MGWA) was performed to predict bacterial genes that influence persistence. In order to perform the MGWA, amino acid sequences were obtained from GenBank for the exact strains we phenotyped. The amino acid sequences of 55 bacterial genomes (Table S7) were clustered into orthologous groups (OGs) using OrthoMCL (37) with an inflation factor of 1.5. The MGWA was then performed using the R package, MAGNAMWAR (38). The inputs for MAGNAMWAR were the clusters of orthologous groups assignments and the CFU per fly at the end of the persistence assay. The MGWA associated OG presence-absence patterns with bacterial persistence levels using a Wilcoxon test. Resultant p-values were Bonferroni corrected and we set an arbitrary significance threshold of p < 0.01. A KEGG enrichment analysis was then done to find functional categories enriched among the significant OGs. BlastKOALA (39) used to assign KEGG functions to a representative sequence from each OG. Pathway significance was then determined by an FDR-corrected chi-square test.

### Motility assay

A motility assay was performed by placing 1μL of OD_600_ normalized bacteria in PBS on mMRS plate with 2 g of agar per liter, replacing the normal plates. The plates were then left at room temperature for 48-72 hours and the diameter of the halo that forms on the plates was measured using a ruler. The tests were performed on flagellar mutants, urea carboxylase mutants, and the wild-type strain *A. fabarum* (Table 2) (40). A linear mixed-effects model was used to test if there was a significant effect of bacterial genotype on the bacterial halo size, and a post-hoc Dunnett test was then applied to determine if there were significant differences between the mean diameter of each mutant versus the wild-type strain.

## DATA AVAILABILITY

All raw data corresponding to the manuscript are included as supporting tables, and are cited throughout the text accordingly.

## ACKNOWLEDGEMENTS

Research reported in this publication was supported in part by the National Institute Of General Medical Sciences of the National Institutes of Health under Award Number R15GM140388. The content is solely the responsibility of the authors and does not necessarily represent the official views of the National Institutes of Health.

## SUPPLEMENTAL FIGURES

**Supplemental Figure 5.**
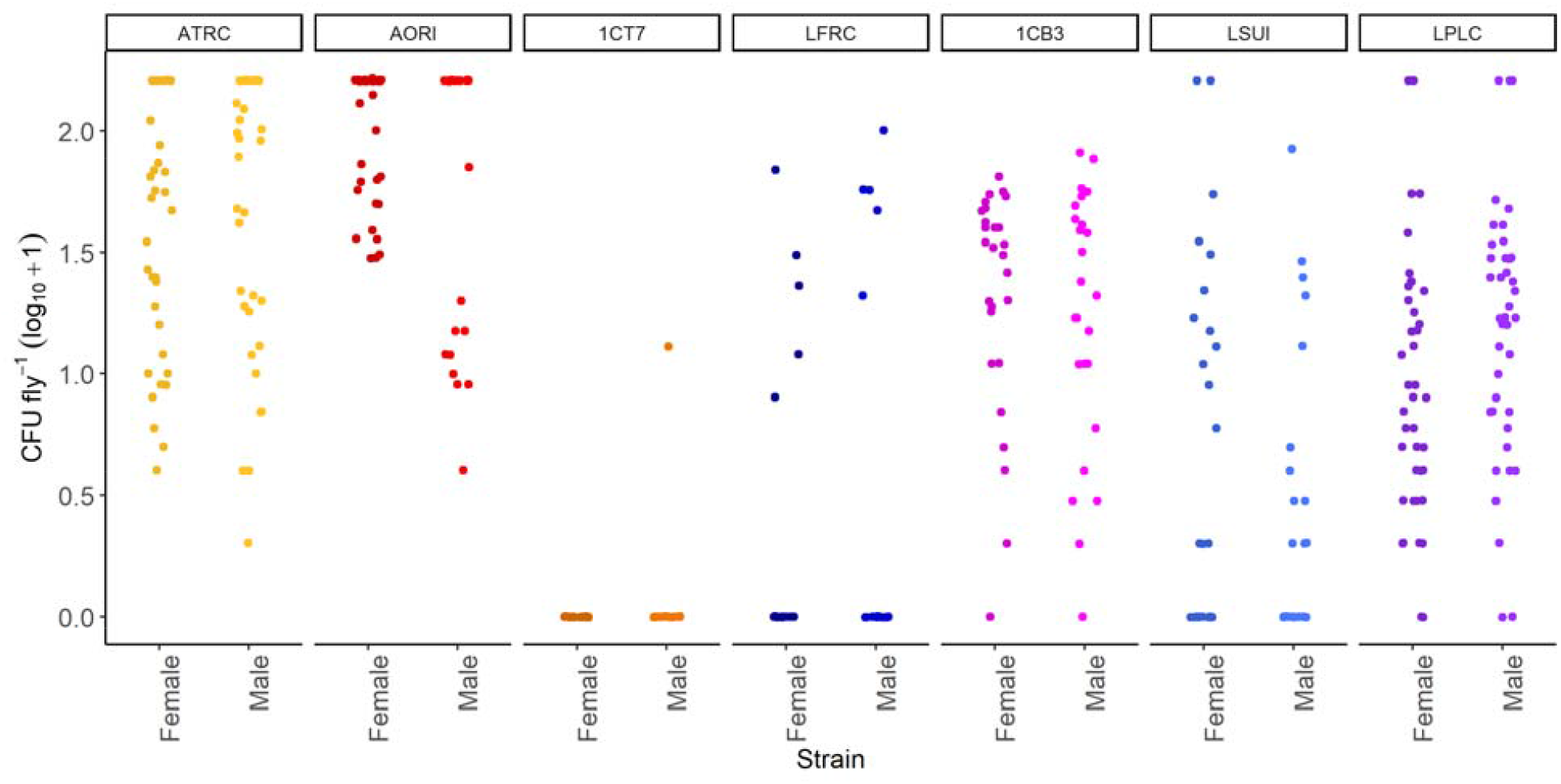
Individual data points show bacterial persistence with the flies is strain-specific. This plot shows the difference in CFU abundances by bacterial strain and fly sex just as Figure 1, but shows each individual data point. Table 3 reports the strain names of the 4-character codes.

**Supplemental Figure 6.**
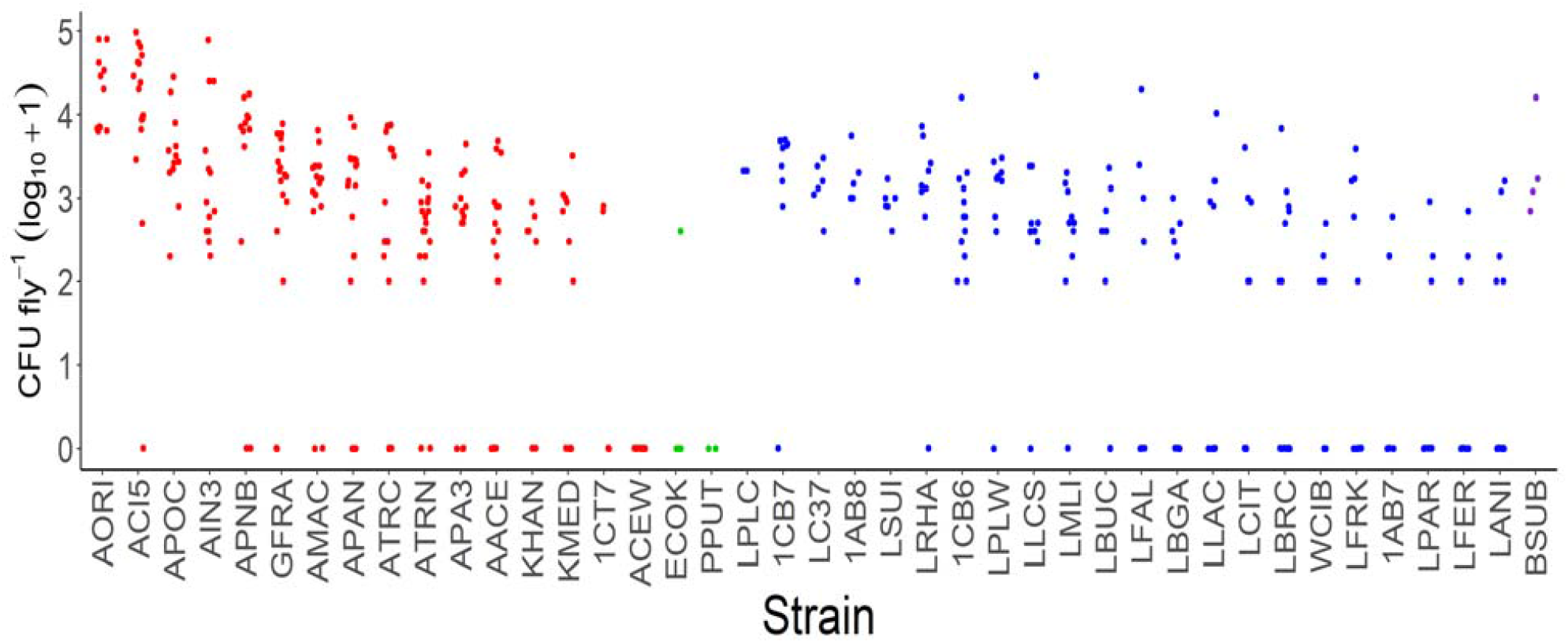
Individual data points show strain-specific differences in bacterial persistence of 41 different bacterial strains. This bar plot shows the log_10_+1 transformed CFU abundances per female fly of 41 different bacterial strains just as Figure 2, but shows each individual data point. Shading matches bacterial groups: acetic acid bacteria (red), gammaproteobacteria (green), lactic acid bacteria (blue), non-lactic acid Firmicutes (purple. Table 3 reports the strain names of the 4-character codes.

**Supplemental Figure 7.**
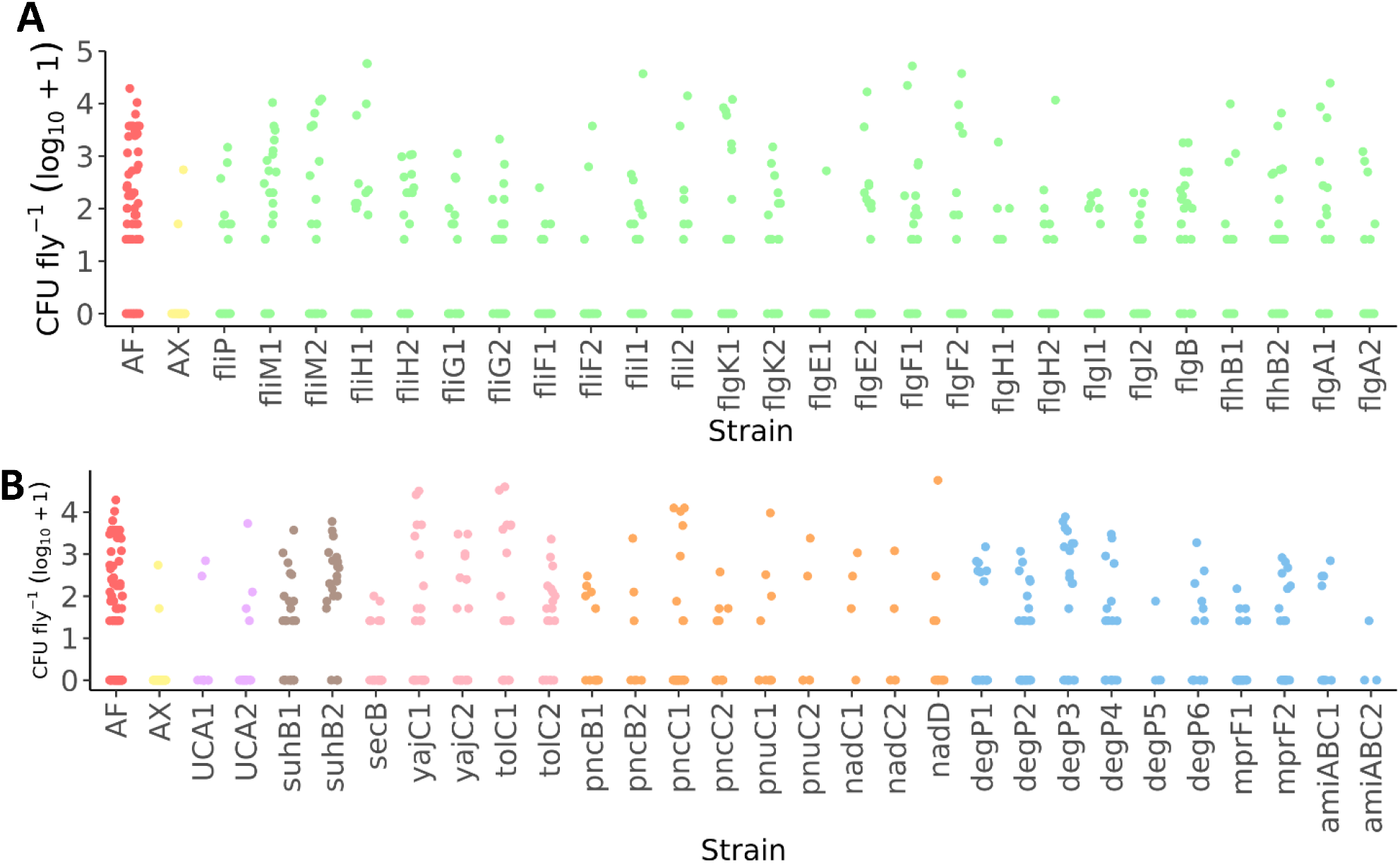
Individual data points show flagellar mutants affect the persistence phenotype. This bar plot shows the log_10_+1 transformed CFU abundances per fly of the mutants tested in the persistence assay just as Figure 3, but with individual data points (A) All non-lagellar mutants testing in the assay. Shading matches type of mutants tested: wild-type *A. fabarum* (red), axenic or bacteria-free flies (yellow), urea carboxylase (purple), phosphatidylinositol (brown), bacterial secretion (pink), nicotinate metabolism (orange), AMP resistance (blue). (B) All flagellar mutants tested in the assay, shown in green. Strain codes are shown in Table 2.

## Notes

### Competing Interest Statement

The authors have declared no competing interest.

